# Shared developmental gait disruptions across two mouse models of neurodevelopmental disorders

**DOI:** 10.1101/2020.10.12.336586

**Authors:** Rachel M. Rahn, Claire T. Weichselbaum, David H. Gutmann, Joseph D. Dougherty, Susan E. Maloney

**Author notes:** Authors contributed equally (Joseph D. Dougherty, Susan E. Maloney). **Correspondence:** Dr. Susan E. Maloney, Washington University School of Medicine, Department of Psychiatry, Campus Box 8232, 660 South Euclid Avenue, St. Louis, MO 63110-1093.

## Abstract

Motor deficits such as abnormal gait are an underappreciated yet characteristic phenotype of many neurodevelopmental disorders (NDDs), including Williams Syndrome (WS) and Neurofibromatosis Type 1 (NF1). Compared to cognitive phenotypes, gait phenotypes are readily and comparably assessed in both humans and model organisms, and are controlled by well-defined CNS circuits. Discovery of a common gait phenotype between NDDs might suggest shared cellular and molecular deficits and highlight simple outcome variables to potentially quantify longitudinal treatment efficacy in NDDs. We therefore characterized gait using the DigiGait assay in two different NDD models: the complete deletion (CD) mouse, which models hemizygous loss of the complete WS locus, and the *Nf1*^+/R681X^ mouse, which models a patient-derived heterozygous germline *NF1* mutation. We collected longitudinal data across five developmental time points (postnatal days 21-30) and one early adulthood time point. Compared to wild type littermate controls, both models displayed markedly similar spatial, temporal, and postural gait abnormalities during development. Developing CD mice also displayed significant decreases in variability metrics. Multiple gait abnormalities observed across *Nf1*^+/R681X^ mouse development persisted into early adulthood, including increased stride length and decreased stride frequency, while developmental abnormalities in CD mice largely resolved by adulthood. These findings suggest that gait subcomponents affected in NDDs show overlap between disorders as well as some disorder-specific features, which may change over the course of development. Our incorporation of spatial, temporal, and postural gait measures also provides a template for gait characterization in other NDD models, and a platform to examining circuits or longitudinal therapeutics.

**Lay Summary:** Gait changes have been reported in Williams Syndrome and Neurofibromatosis Type 1, but how these changes develop over time has not been explored. We therefore studied gait in mouse models of these two disorders across time. We found multiple shared differences in gait as compared to healthy controls at the younger ages in both models. However, those differences were resolved in the Williams Syndrome model by adulthood, yet persisted in the Neurofibromatosis Type 1 model.

## INTRODUCTION

Despite a growing understanding of the clinical deficits that characterize neurodevelopmental disorders (NDDs), motor impairments remain an understudied yet prevalent phenotype across many NDDs. Motor phenotypes such as abnormal gait are a well-documented feature pervasive in NDDs (Hilton et al., 2012; Hocking et al., 2009; Kindregan et al., 2015; Kraan et al., 2017; Sanders & Gillig, 2010) and represent a brain function and circuit better understood and more highly conserved across humans and mice than that of the social and cognitive phenotypes also characteristic to NDDs. Thus, they represent an opportunity to define the consequences of NDD-associated genetic mutations on the development and function of well-understood CNS circuits, and to determine whether distinct neurogenetic disorders might have some shared consequences. Furthermore, these more accessible phenotypes could be used to judge both the efficacy of NDD treatments as well as the effect of long-term use of these therapeutics. However, the existence of a common gait phenotype between multiple NDDs has not been established. This is due to the difficulty of performing longitudinal human studies and the confounding effects of genetic heterogeneity in clinical populations.

Williams Syndrome (WS) and Neurofibromatosis Type 1 (NF1) are two NDDs with known gait disruptions and well-understood genetic etiologies, making them ideal options for a focused examination of a subsection of the heterogeneous NDD clinical population. Gait abnormalities in WS are known to occur during childhood and persist into adulthood (Hocking et al., 2009; Morris et al., 2020). NF1 gait phenotypes have also been reported during childhood and adolescence (Champion et al., 2014), but their trajectory into adulthood is less well-documented. In addition, gait abnormalities are among the motor phenotypes observed in approximately 80% of autism spectrum disorder (ASD) cases (Hilton et al., 2012), and almost half of all NF1 patients exhibit at least partial ASD symptomatology (Garg et al., 2013; Morris et al., 2016). Specifically, about 25% of NF1 patients exhibit a phenotypic profile consistent with idiopathic ASD, and another 20% exhibit partial ASD symptomatology (Garg et al., 2013; Morris et al., 2016). Recently, the *NF1* gene was confirmed as a quantitative trait loci for ASD (Morris et al., 2016). This large-scale study identified that the Quantitative Autistic Trait (QAT) scores for NF1 individuals were continuously distributed and pathologically shifted. Whether these NF1 gait phenotypes are a result of ASD overlap or are an independent feature of NF1 has yet to be determined.

Mouse models provide an excellent opportunity to directly target the specific effects of genotype on phenotype in NDDs and to map the neural circuitry underlying abnormalities in these conditions. Production of complex motor behavior, such as gait, is a conserved behavior and robustly studied in mice and humans. Besides allowing for the study of targeted alleles and providing brain samples to understand cellular and molecular consequences, mouse models also permit longitudinal studies to be performed in a much shorter timeframe than in humans, so that subtle changes in developmental trajectories across the population can be investigated. Crucially, studying multiple mouse models of NDD permits the execution of well-powered studies capable of identifying phenotypes too subtle to detect in clinical study populations, as well as the investigation of commonalities across models.

For these reasons, we characterized gait during development and in adulthood in mouse models of WS and NF1 in order to determine whether developmental gait abnormalities in these NDDs resolve or persist into adulthood. Using our recently developed DigiGait pipeline (Akula et al., 2020), we quantified gait in the Complete Deletion (CD) mouse model hemizygous for the 26 gene WS locus (Segura-Puimedon et al., 2014) and a mouse model harboring a patient-derived heterozygous loss-of-function missense mutation in the murine homologue of the human *NF1* gene (*Nf1*^+/R681X^) (Li et al., 2016; Pinna et al., 2015; Rojnueangnit et al., 2015). We found that both the CD and *Nf1*^+/R681X^ models exhibited gait alterations during development relative to WT littermates, many of which persisted into adulthood in the *Nf1*^+/R681X^ mice but not the CD mice. Unexpectedly, across these two different models of NDD, the abnormalities during development were nearly identical. These results suggest that these NDDs share a common gait phenotype and provide a foundation for the future study of treatments that may rescue motor circuit dysfunction in these models.

## METHODS

### Animals

All experimental protocols were approved by and performed in accordance with the relevant guidelines and regulations of the Institutional Animal Care and Use Committee of Washington University in St. Louis, and were in compliance with US National Research Council’s Guide for the Care and Use of Laboratory Animals, the US Public Health Service’s Policy on Humane Care and Use of Laboratory Animals, and Guide for the Care and Use of Laboratory Animals. All mice (Mus musculus) used in this study were maintained and bred in the vivarium at Washington University in St. Louis. The colony room lighting was 12:12h light/dark cycle; room temperature (~20-22°C) and relative humidity (50%) controlled automatically. Standard lab diet and water were freely available. Pregnant dams were individually housed in translucent plastic cages measuring 28.5×17.5×12cm with corncob bedding. Upon weaning at postnatal day (P)21, mice for behavioral testing were group housed according to sex.

Gait was assessed in two mouse lines modeling genetic risk for the neurodevelopmental disorders Neurofibromatosis Type 1 (NF1) and Williams Syndrome (WS). The NF1 model harbored a NF1-patient-derived *Nf1* gene mutation (c.2041C>T;p.R681X; *Nf1*^+/R681X^) (Toonen et al., 2016). To generate this cohort, *Nf1*^+/R681X^ mice were crossed to C57BL/6J wild type (WT; RRID:IMSR_JAX:000664) mice to produce 30 *Nf1*^+/R681X^ (14 males, 16 females) and 40 WT (24 males, 16 females) littermates. Homozygous *Nf1* deletion is embryonic lethal and not found in patients with NF1 (Gutmann & Giovannini, 2002), and thus was not modeled here. The WS model harbors a heterozygous complete deletion (CD) of the WS critical region comprising 26 genes on the conserved syntenic region of chromosome 5 in the mouse (Segura-Puimedon et al., 2014). This cohort was generated by crossing CD mice to WT mice to generate 24 CD (9 males, 15 females) and 44 WT (19 males, 25 females) littermates. Homozygous deletion of the WS critical region also is embryonic lethal.

### Gait Analysis

Gait data was collected using the DigiGait Imaging System (Mouse Specifics, Inc.; Framingham, MA), an advanced gait analysis system with Ventral Plane Imaging Technology that generates digital paw prints from the animal as it runs on a motorized treadmill (Hampton et al., 2004). All behavioral experiments were performed by a female experimenter blinded to genotype. All behavioral testing occurred during the light phase between 12:00pm – 6:00pm. Gait performance was collected four times across development and again during adulthood (Figure 1A) as previously described (Akula et al., 2020). Briefly, mice were habituated to the apparatus environment and belt movement at P20. Gait testing occurred at P21, P24, P27, P30 and once again after P60. On each test day, a video was captured of each animal’s gait at an age-appropriate belt speed (20 cm/s during development and 30 cm/s in adulthood). Gait video processing and metric selection, as well as body length quantification, were conducted as previously described (Akula et al., 2020).

**Figure 1.**
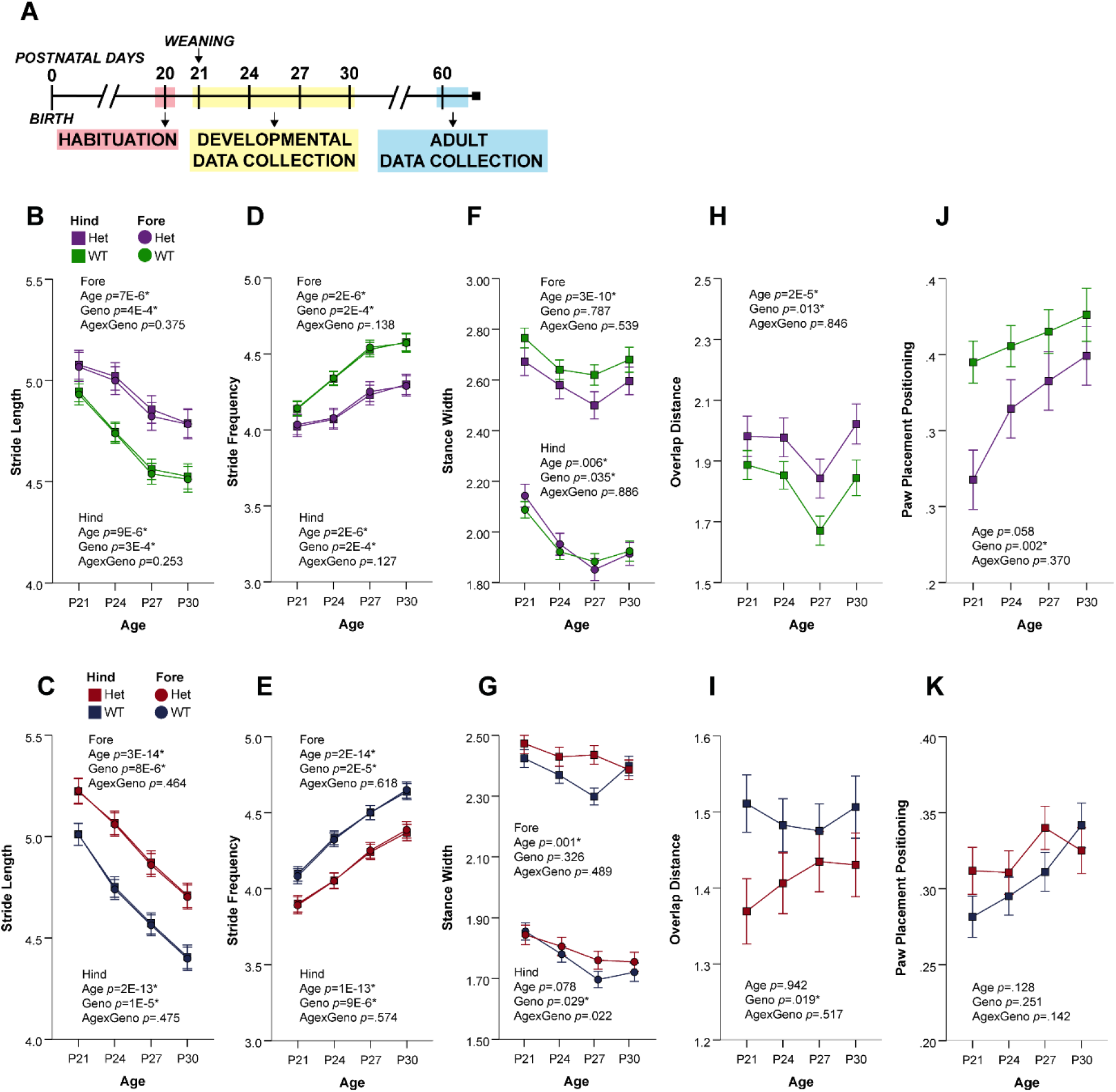
Developing CD and *Nf1*^+/R681X^ mice significantly differ from controls in spatial gait subcomponents. **A)** Timeline of DigiGait data collection schedule. **B-K)** Change in forelimb and hindlimb metrics of B,C) stride length, D,E) stride frequency, F,G) stance width, H,I) overlap distance, and J,K) paw placement positioning at P21, P24, P27, and P30 for CD mice (purple=het, N=24; green=WT, N=44), and for *Nf1*^+/R681X^ mice (red=het, N=30; blue=WT, N=40). Data are means ±SEM. * indicates significance after FDR correction. Hindlimb means are represented by squares, and forelimb means are represented by circles.

### Statistical Analysis

All statistical analyses were performed using IBM SPSS Statistics software (v.25, RRID:SCR_002865). Prior to analyses, all data were screened for missing values, fit of distributions with assumptions underlying univariate analyses, and influential outliers. Hierarchical linear mixed modeling was used to analyze gait data across juvenile ages, with genotype, sex and age as fixed effects. Age was also used as a repeated random factor clustered by subject ID. Body length was added as a covariate to account for the influence on gait of changing body length across development. Adult data was analyzed using a two-way ANCOVA, with genotype and sex as fixed effects and body length as a covariate. The Benjamini-Hochberg correction for False Discovery Rate (FDR; at *q*=.1) was used to adjust the critical alpha level for multiple analyses within each mouse line. A fixed effect of sex was not observed in any gait parameter in either model reported here, except for stance width variability during development in *Nf1*^+/R681X^ males. Therefore, all data below, except for *Nf1*^+/R681X^ forelimb stance width variability, is collapsed across sex. The datasets generated and analyzed in this study are available from the corresponding author upon reasonable request.

## RESULTS

Cross-sectional studies of gait in NDD models reveal disorder symptomatology at a particular point in time, yet they cannot determine whether gait abnormalities reflect a delay in typical development or a persistent abnormality. Resolution of gait defects by adulthood would suggest developmental delay, while persistence in adulthood would reflect a more permanent phenotype. Therefore, we used a longitudinal design to quantify gait across development and in adulthood to parse resolution versus persistence of gait phenotypes in *Nf1*^+/R681X^ and WS mouse models, as well as characterize a comprehensive set of spatial, temporal, and postural gait subcomponents and identify those most affected in these NDDs.

### Trajectory of gait development over time replicates previous findings independent of genotype

Previous studies have identified multiple spatial, temporal, and postural characteristics of gait that vary across development in the C57Bl/6J background strain used here (Akula et al., 2020). When collapsed across genotype, both models in the present study largely replicated this previous work; no significant age by genotype interactions were observed. As previously reported, spatial subcomponents such as stance width, excepting the *Nf1*^+/R681X^ hindlimb measure, varied significantly across age when compared to WT littermates in both the *Nf1*^+/R681X^ and CD mice. Temporal variables such as swing and stance duration in both fore- and hindlimbs and hindlimb propulsion duration also changed in our models across age as in the previous study, as well as postural elements such as peak paw area and maximum rate of paw contact change of the hindpaws (Table 1) (Akula et al., 2020). Some additional age-dependent effects independent of genotype not previously reported, such as changes in stride frequency, were also observed. Overall, these findings confirmed that the progression of baseline, healthy gait development across age in our controls aligned with the age-dependent gait changes previously reported, and confirm that our genotype-dependent findings were based in disease-related dysfunction.

**Table 1.**
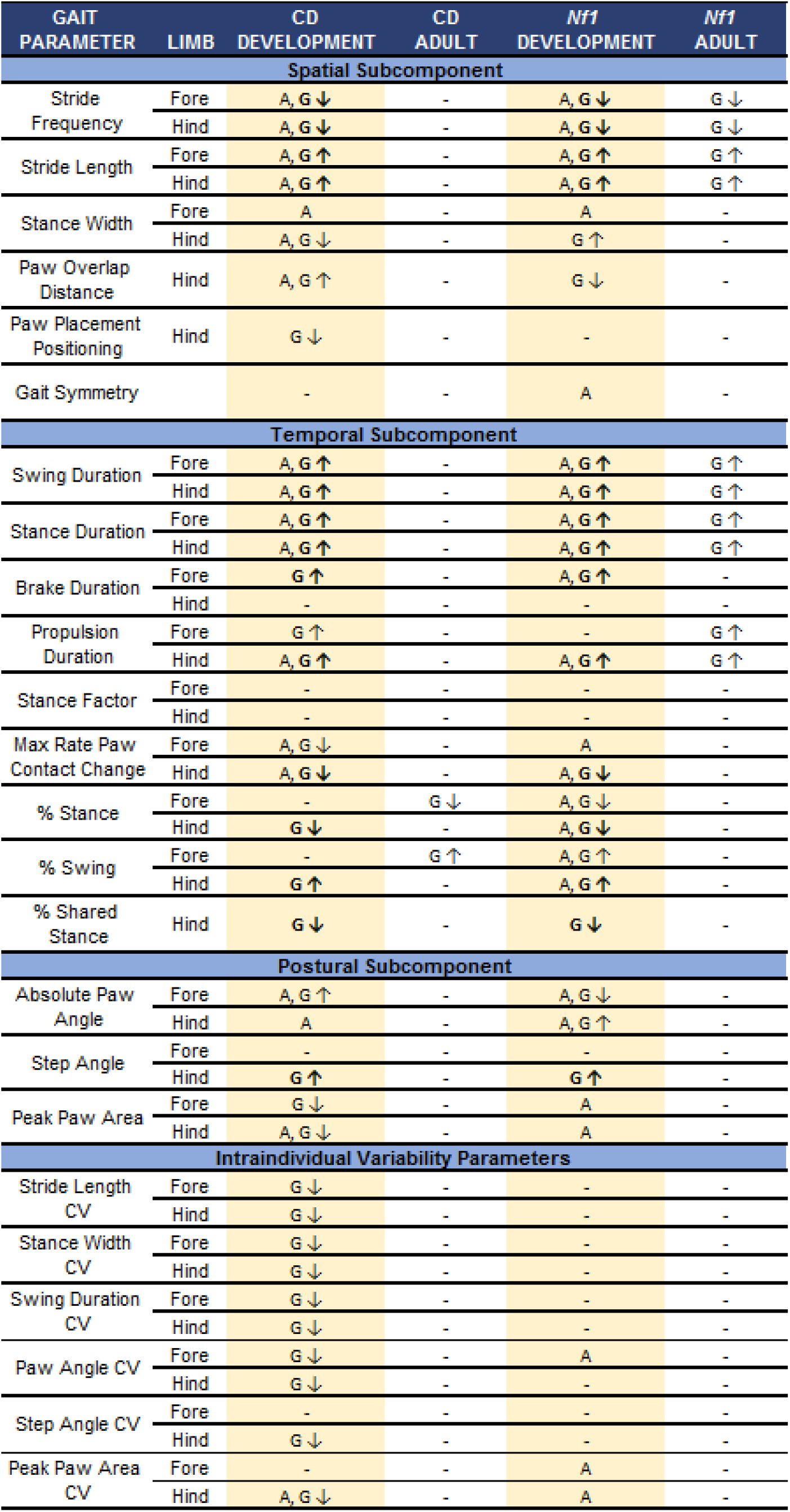
Summary of significant findings for juvenile mice and adult mice across both *Nf1* and CD models. For each gait parameter, main effects of age (A) and genotype (G) are indicated where significant after FDR correction. Genotype effects shared across models are in bold. Age by genotype interactions were also explored but none survived correction. Arrows indicate the direction of the genotype effect regarding Het compared to WT (down arrows indicate lower score for Het compared to WT; up arrows indicate higher score for Het compared to WT). N = 24 Het CD mice and 44 WT CD mice; N = 30 Het *Nf1*^+/R681X^ mice and 40 WT *Nf1*^+/R681X^ mice.

| GAIT<br>PARAMETER | LIMB | CD<br>DEVELOPMENT | CD<br>ADULT | *Nf1*<br>DEVELOPMENT | *Nf1*<br>ADULT |
| --- | --- | --- | --- | --- | --- |
| Spatial Subcomponent |  |  |  |  |  |
| Stride<br>Frequency | Fore | A, G ↓ | - | A, G ↓ | G ↓ |
|  | Hind | A, G ↓ | - | A, G ↓ | G ↓ |
| Stride Length | Fore | A, G ↑ | - | A, G ↑ | G ↑ |
|  | Hind | A, G ↑ | - | A, G ↑ | G ↑ |
| Stance Width | Fore | A | - | A | - |
|  | Hind | A, G ↓ | - | G ↑ | - |
| Paw Overlap<br>Distance | Hind | A, G ↑ | - | G ↓ | - |
| Paw Placement<br>Positioning | Hind | G ↓ | - | - | - |
| Gait Symmetry |  | - | - | A | - |
| Temporal Subcomponent |  |  |  |  |  |
| Swing Duration | Fore | A, G ↑ | - | A, G ↑ | G ↑ |
|  | Hind | A, G ↑ | - | A, G ↑ | G ↑ |
| Stance Duration | Fore | A, G ↑ | - | A, G ↑ | G ↑ |
|  | Hind | A, G ↑ | - | A, G ↑ | G ↑ |
| Brake Duration | Fore | G ↑ | - | A, G ↑ | - |
|  | Hind | - | - | - | - |
| Propulsion<br>Duration | Fore | G ↑ | - | - | G ↑ |
|  | Hind | A, G ↑ | - | A, G ↑ | G ↑ |
| Stance Factor | Fore | - | - | - | - |
|  | Hind | - | - | - | - |
| Max Rate Paw<br>Contact Change | Fore | A, G ↓ | - | A | - |
|  | Hind | A, G ↓ | - | A, G ↓ | - |
| % Stance | Fore | - | G ↓ | A, G ↓ | - |
|  | Hind | G ↓ | - | A, G ↓ | - |
| % Swing | Fore | - | G ↑ | A, G ↑ | - |
|  | Hind | G ↑ | - | A, G ↑ | - |
| % Shared<br>Stance | Hind | G ↓ | - | G ↓ | - |
| Postural Subcomponent |  |  |  |  |  |
| Absolute Paw<br>Angle | Fore | A, G ↑ | - | A, G ↓ | - |
|  | Hind | A | - | A, G ↑ | - |
| Step Angle | Fore | - | - | - | - |
|  | Hind | G ↑ | - | G ↑ | - |
| Peak Paw Area | Fore | G ↓ | - | A | - |
|  | Hind | A, G ↓ | - | A | - |
| Intraindividual Variability Parameters |  |  |  |  |  |
| Stride Length<br>CV | Fore | G ↓ | - | - | - |
|  | Hind | G ↓ | - | - | - |
| Stance Width<br>CV | Fore | G ↓ | - | - | - |
|  | Hind | G ↓ | - | - | - |
| Swing Duration<br>CV | Fore | G ↓ | - | - | - |
|  | Hind | G ↓ | - | - | - |
| Paw Angle CV | Fore | G ↓ | - | A | - |
|  | Hind | G ↓ | - | - | - |
| Step Angle CV | Fore | - | - | - | - |
|  | Hind | G ↓ | - | - | - |
| Peak Paw Area<br>CV | Fore | - | - | A | - |
|  | Hind | A, G ↓ | - | A | - |

### *Both* Nf1^*+/R681X*^ and CD models display abnormal spatial, temporal, and postural gait characteristics in development

Consistent with NDD risk factors disrupting gross motor development, both *Nf1*^+/R681X^ and CD models displayed significance differences compared to WT littermate controls in multiple spatial, temporal, and postural subcomponents of gait during development (Table 1). Stride length, an important spatial gait subcomponent, was significantly increased in both models as compared to their WT littermate controls (Figure 1B,C). Correspondingly, stride frequency was significantly decreased in both CD mice and *Nf1*^+/R681X^ mice relative to controls (Figure 1D,E). Spatial gait abnormalities in stance width and overlap distance were also observed in both models, with a narrower hindlimb stance width for CD mice than controls and a wider hindlimb stance width in *Nf1*^+/R681X^ mice (Figure 1F,G). There was also an increased overlap distance in CD mice, which was decreased in *Nf1*^+/R681X^ mice compared to controls (Figure 1H,I). The only gait component that displayed a sex effect in either model was *Nf1*^+/R681X^ forelimb stance width coefficient of variance (CV), a measure of relative variability (females 14.562±.417, males 13.278±.385), but an interaction between sex and genotype was not a predictor of outcome. Furthermore, genotype was not a significant predictor of forelimb stance width CV for either females or males when analyzed separately. In addition, hindpaw placement positioning was significantly reduced in the CD but not *Nf1*^+/R681X^ mice as compared to WT littermates (Figure 1J,K), suggesting that spatial gait abnormalities are not entirely identical between these two NDD models despite their remarkable similarities.

In terms of temporal subcomponents of the gait phenotype, both CD and *Nf1*^+/R681X^ models displayed significant changes in swing duration and other related metrics as compared to WT littermates. In both fore- and hindlimb, swing duration was significantly increased in developing CD mice and *Nf1*^+/R681X^ mice when compared to WT controls (Figure 2A,B). The percentage of the total stride cycle that was spent in the stance phase was significantly decreased in CD hindlimbs as well as both *Nf1*^+/R681X^ fore- and hindlimbs, while propulsion duration was longer in CD fore- and hindlimbs and *Nf1*^+/R681X^ hindlimbs compared to WT littermates (Figure 2C-F).

**Figure 2.**
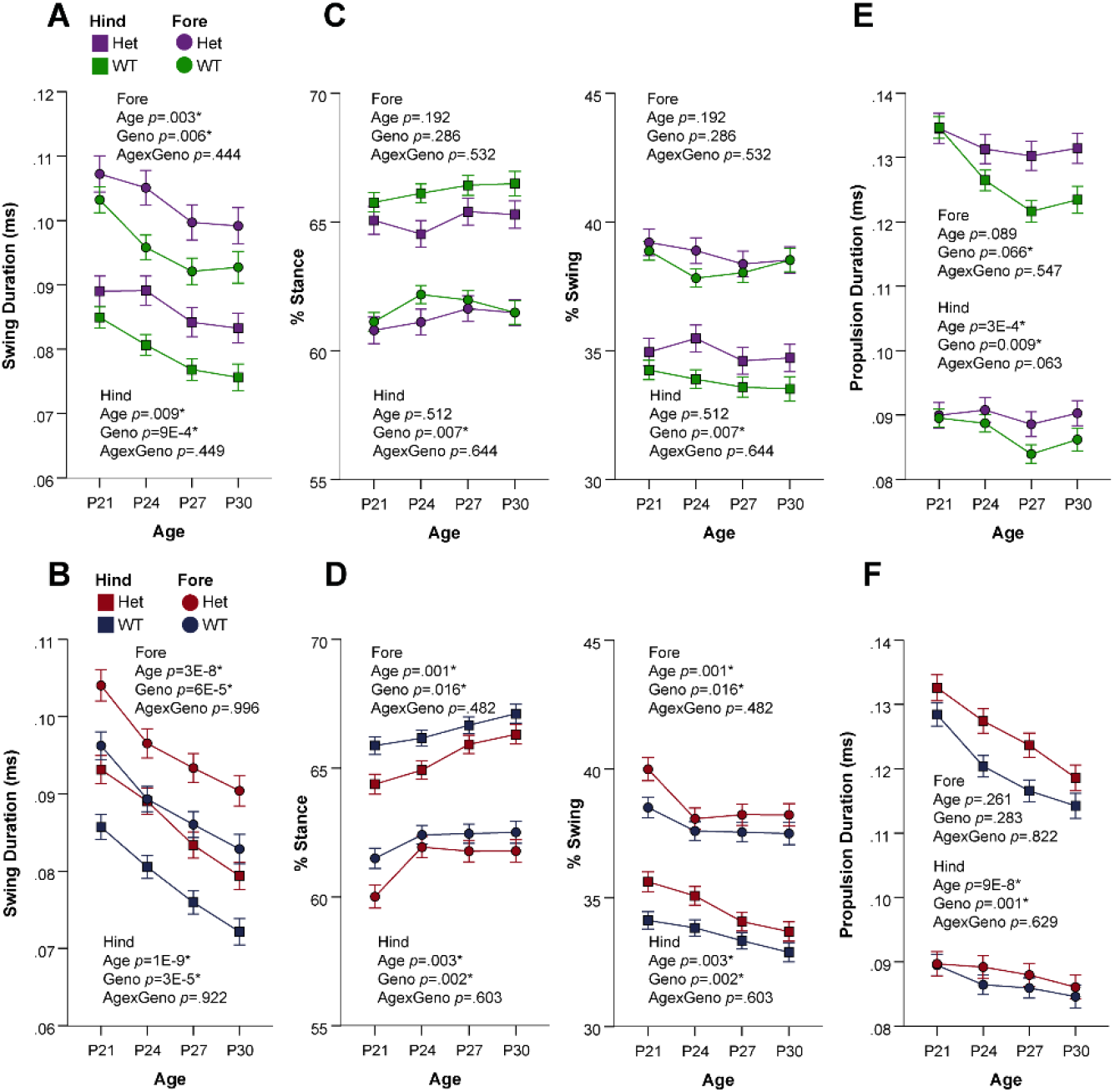
Developing CD and *Nf1*^+/R681X^ mice significantly differ from controls in temporal gait subcomponents. Change in forelimb and hindlimb metrics of A,B) swing duration, C,D) percentage of stride cycle that is stance phase (left) and swing phase (right), and E,F) propulsion duration at P21, P24, P27, and P30 for CD mice (purple=het, N=24; green=WT, N=44) and for *Nf1*^+/R681X^ mice (red=het, N=30; blue=WT, N=40). Data are means ±SEM. * indicates significance after FDR correction. Hindlimb means are represented by squares, and forelimb means are represented by circles.

In addition to spatial and temporal characteristics of gait, postural differences in both NDD models were evident when compared to controls. The absolute angle of the forepaws was significantly increased in both CD and *Nf1*^+/R681X^ models (Figure 3A,B), suggesting that these animals show an altered body positioning while walking. While *Nf1*^+/R681X^ mice did not display any other significant abnormalities in postural gait metrics, CD animals showed a significant reduction in peak paw area on the walking surface for both fore- and hindpaws (Figure 3C,D). These findings suggest that posture while walking is significantly impacted in development in both the *Nf1*^+/R681X^ and CD models.

**Figure 3.**
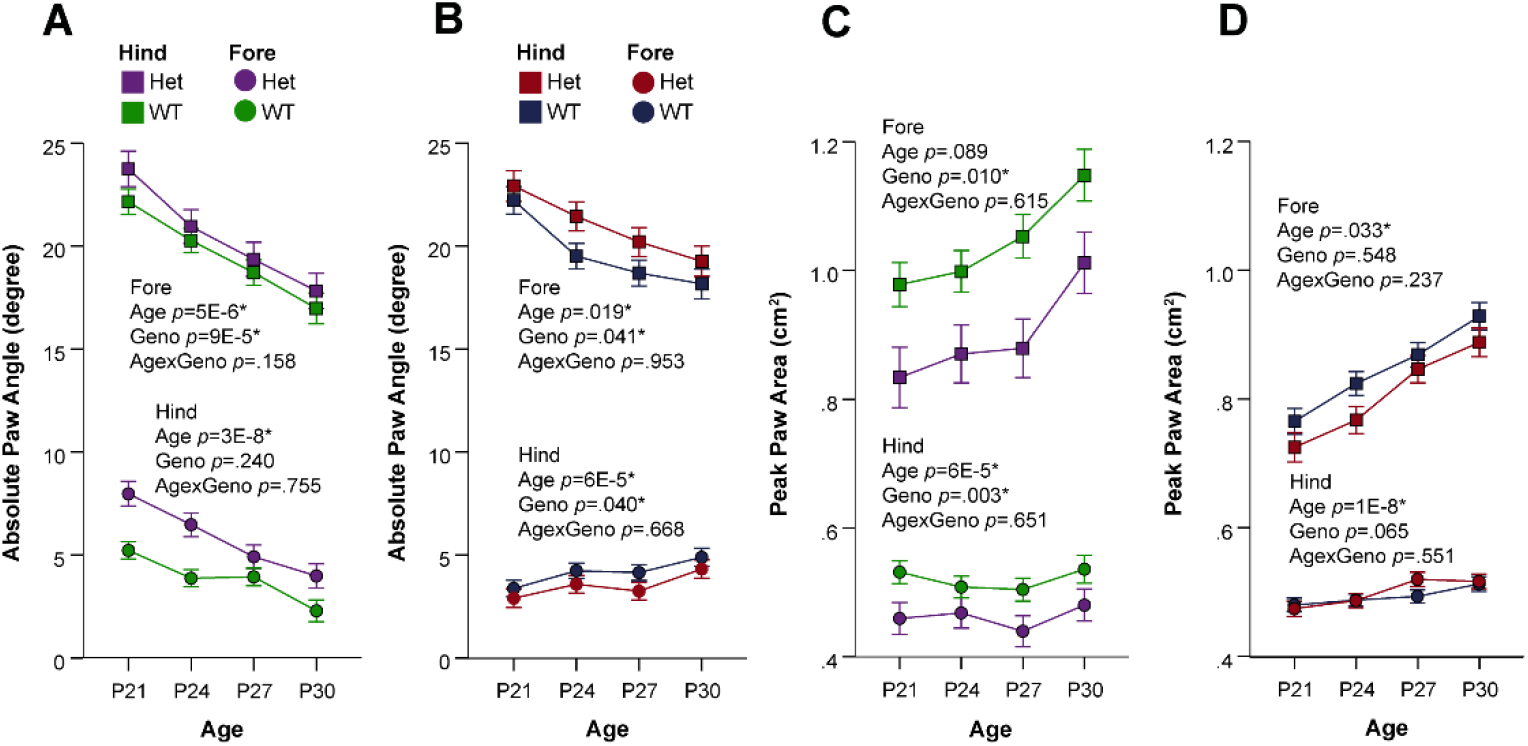
Developing CD and *Nf1*^+/R681X^ mice significantly differ from controls in postural gait subcomponents. Change in forelimb and hindlimb metrics of A,B) absolute paw angle and C,D) peak paw angle at P21, P24, P27, and P30 for CD mice (purple=het, N=24; green=WT, N=44) and for *Nf1*^+/R681X^ mice (red=het, N=30; blue=WT, N=40). Data are means ±SEM. * indicates significance after FDR correction. Hindlimb means are represented by squares, and forelimb means are represented by circles.

### Gait abnormalities persist into early adulthood for Nf1^+/R681X^ mice but largely resolve for CD mice

We have thus far reported our characterization of developmental gait abnormalities in both the CD and *Nf1*^+/R681X^ models. We also quantified gait in these models in adulthood to investigate whether these phenotypes reflect a developmental delay in gait and resolution by adulthood, or reflect a permanent abnormality and persistence into adulthood. In contrast to the multiple spatial, temporal, and postural abnormalities in development, adult CD mice only display one significant difference compared to WT littermates, a decrease in the percentage of the forelimb stride cycle spent in the stance phase and an accompanying increase in the percentage spent in swing (Figure 4A-F). While temporal stance and swing abnormalities were also present in CD mice during development, their decreased stance phase percentage was observed in the hindlimbs, not forelimbs, and therefore the temporal stride cycle phenotype observed in adulthood does not appear to have directly persisted from the developmental period (Table 1). Conversely, *Nf1*^+/R681X^ mice displayed numerous phenotypes that persisted from development into adulthood, including significantly increased stride length and decreased stride frequency, as well as longer stance, swing, and propulsion durations in both fore- and hindlimbs relative to WT littermates (Figure 4G-L). Although approximately half of the gait metrics perturbed in developing *Nf1*^+/R681X^ mice no longer showed significant differences in adulthood, developmental and adult *Nf1*^+/R681X^ datasets displayed a high degree of concordance in the direction of effect for the gait alterations that remained present in adulthood. The perturbed gait variables in *Nf1*^+/R681X^ adults also tended to be the most highly significant variables during development, such as stride length, which was significantly increased in both development and adulthood as compared to WT littermates (Figure 5). Overall, these findings demonstrate that *Nf1*^+/R681X^ mice exhibit multiple spatial, temporal, and postural gait abnormalities in gait subcomponents that persist into adulthood, while the CD model has developmental disruptions in gait that are largely resolved by adulthood.

**Figure 4.**
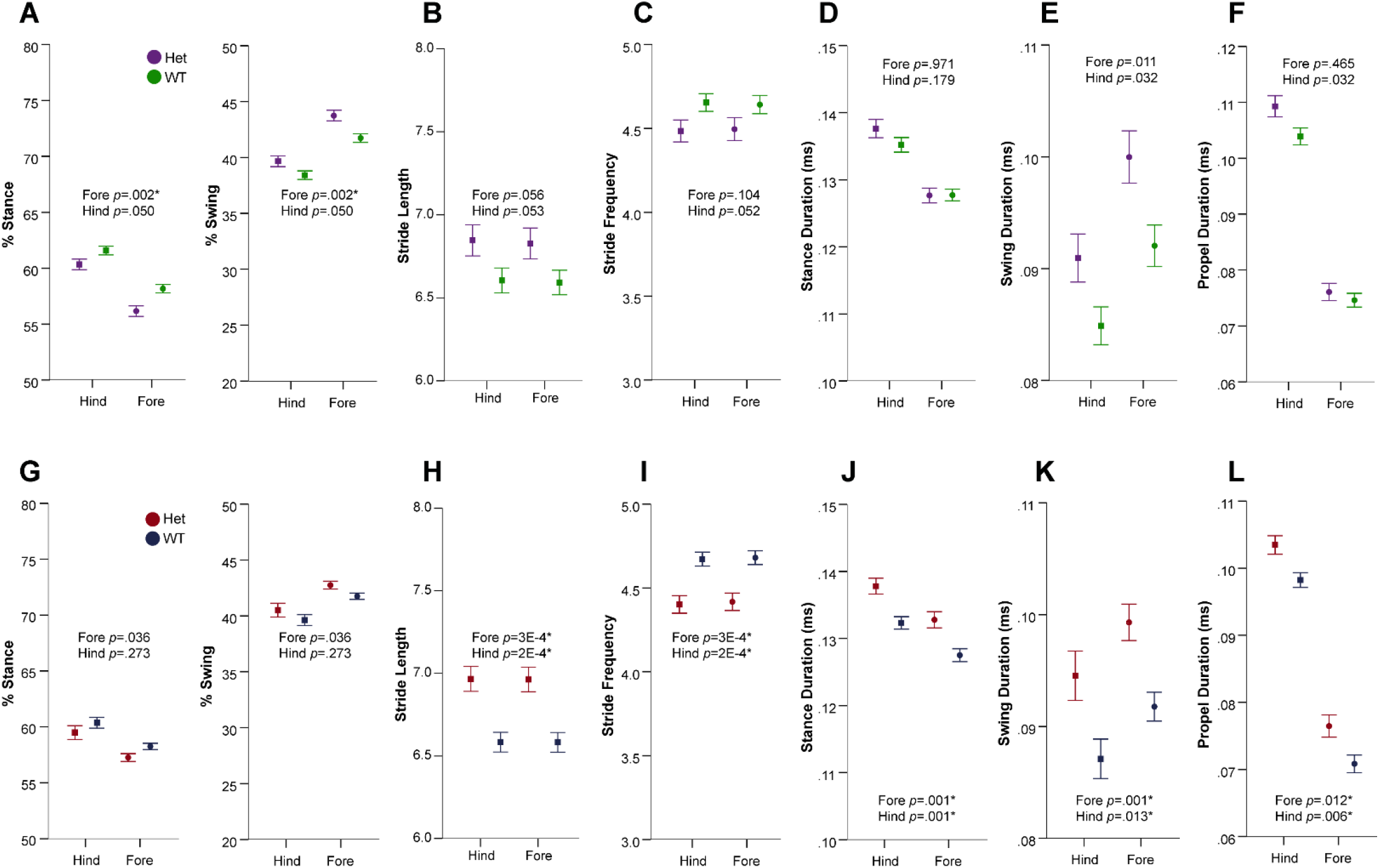
*Nf1*^+/R681X^ mouse gait abnormalities are sustained into early adulthood while CD abnormalities are mostly resolved. **A-F)** Change in forelimb and hindlimb metrics of A,G) percentage of stride cycle that is stance phase (left) and swing phase (right), B,H) stride length, C,I) stride frequency, D,J) stance duration, E,K) swing duration, and F,L) propulsion duration after P60 for CD mice (purple=het, N=24; green=WT, N=44) and for *Nf1*^+/R681X^ mice (red=het, N=30; blue=WT, N=40). Data are means ±SEM. * indicates significance after FDR correction.

**Figure 5.**
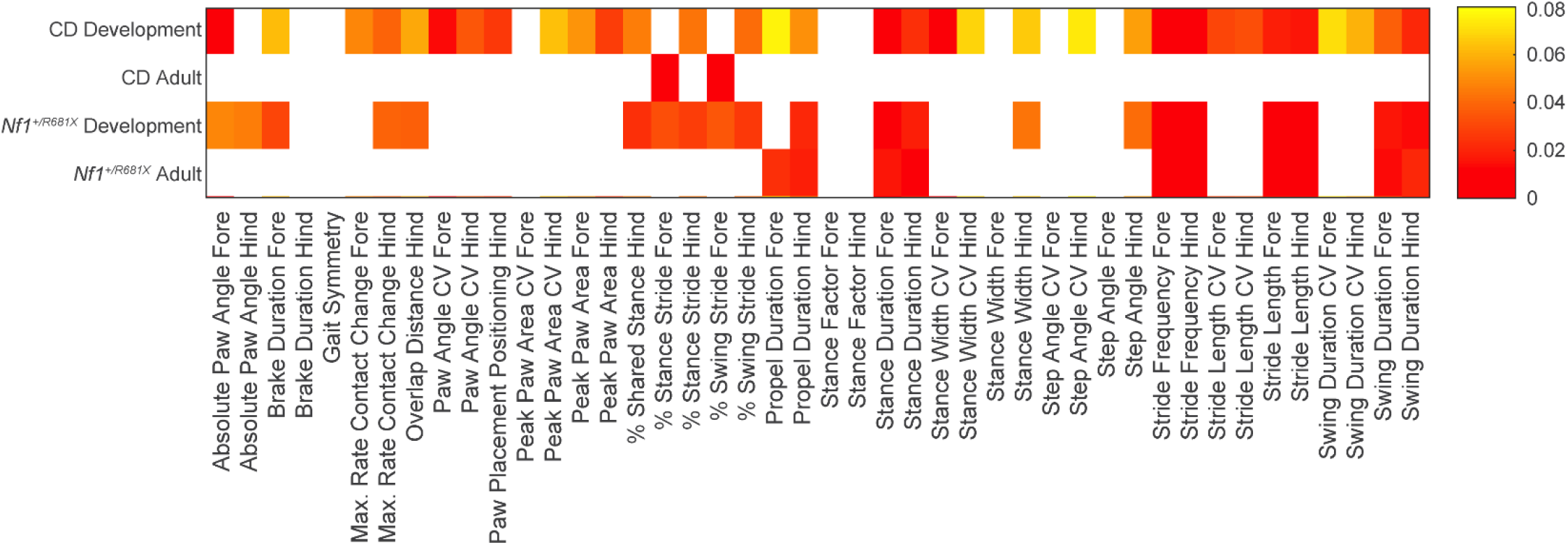
*Nf1*^+/R681X^ and CD mice share multiple significant gait alterations during development but not early adult timepoints. Heat map of the significance level by age of each gait metric from the CD and *Nf1*^+/R681X^ cohorts in development (P21-P30, linear mixed modeling) and adulthood (after P60, ANCOVA). Values represent significant new critical alpha values after Benjamini-Hochberg correction. Smallest values represent metrics with the lowest uncorrected *p* and highest significance ranking of 44 metrics measured. White represents metrics that are not significant after multiple comparison correction.

### Reduced intraindividual gait variability is evident in CD but not Nf1^+/R681X^ mice in development but resolves by adulthood

In both clinical populations as well as mouse models of NDD, a compelling question regarding development remains whether individuals show significant variability in their own gait or other symptomatology. For typically developing children, gait variability decreases across development as children become more proficient at walking and take more consistent steps (Hausdorff et al., 1999). However, children with NDDs often exhibit greater gait variability, including increased variability in stride length and other metrics in children with ASD (Nayate et al., 2012; Rinehart et al., 2006). Adults with WS also show increased intraindividual variability in stride length and other measures (Hocking et al., 2009). When comparing intraindividual variability of *Nf1*^+/R681X^ animals and WT littermates, no significant differences were observed, but CD mice did significantly differ in their variability of several gait metrics compared to their WT littermates. Specifically, variability in multiple gait measures that included stride length, stance width, and swing duration were all reduced in developing CD mice compared to WT littermates (Figure 6, Table 1). This reduced variability did not persist into adulthood, as no gait metrics displayed significant differences in variability for adult CD mice relative to WT littermates (Table 1). This implies that the reduced gait variability observed in the CD model occurs only during development and resolves by adulthood.

**Figure 6.**
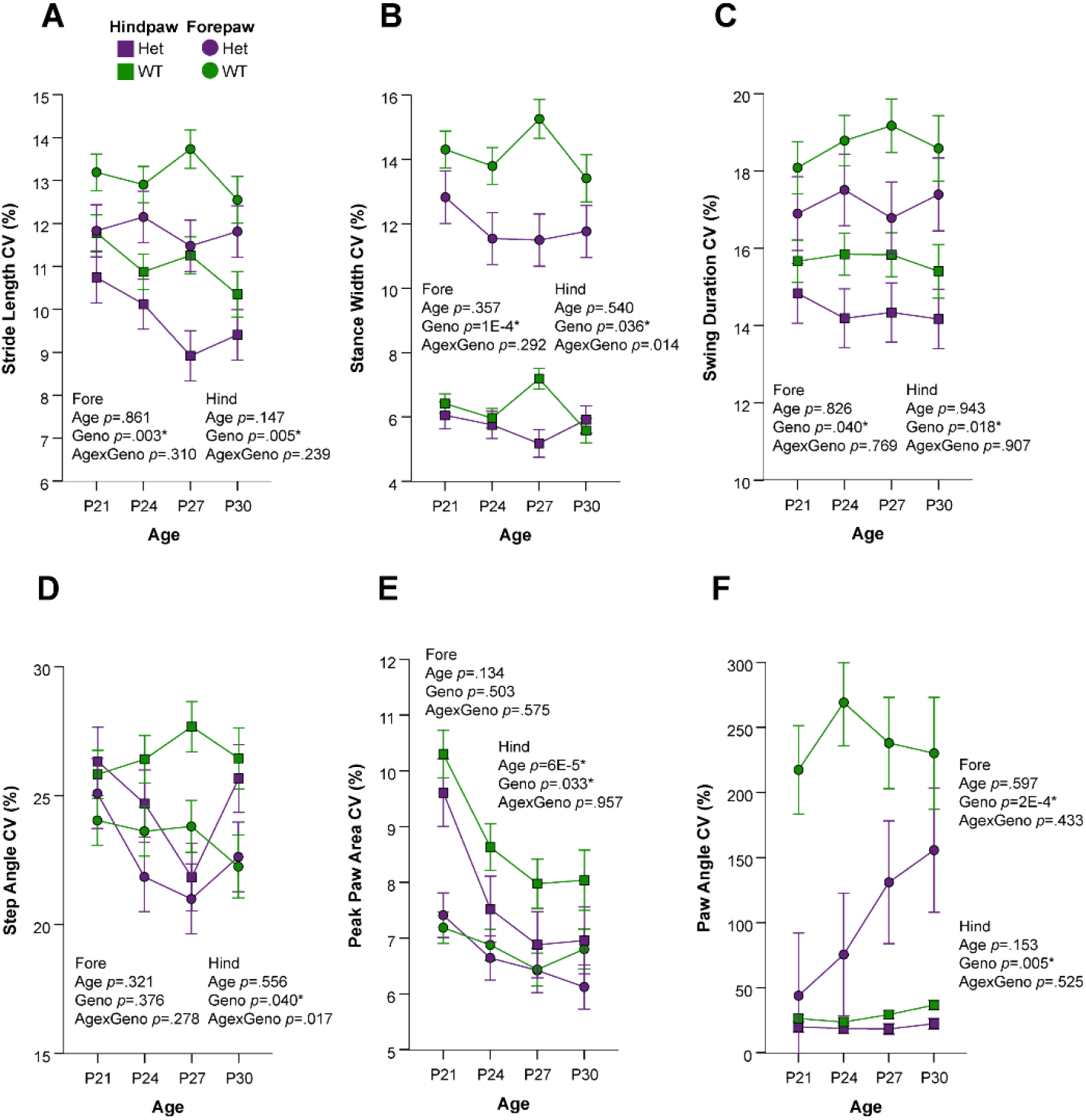
Magnitude of CD developmental gait variability metrics is significantly smaller than controls. Change in CD forelimb and hindlimb coefficient of variance (CV) metrics of A) stride length, B) stance width, C) swing duration, and D) step angle at P21, P24, P27, and P30 (purple=het, N=24; green=WT, N=44). Data are means ±SEM. * indicates significance after FDR correction. Hindlimb means are represented by squares, and forelimb means are represented by circles.

## DISCUSSION

Through characterization of gait during development and adulthood in mouse models of WS and NF1, we identified, for the first time, aspects of gait perturbed by these specific genetic NDD liabilities in a model system. Furthermore, we sought to determine whether developmental gait abnormalities in these NDDs resolve by adulthood and would therefore represent a delay in gait development, rather than a persistent abnormality. Our findings indicated that the *Nf1*^+/R681X^ mouse, which models a patient-derived mutation, and the CD mouse, which models hemizygous loss of the complete WS locus, both exhibit abnormalities in multiple features of gait that are evident during development. These features included an increase in stride length, propulsion duration, and swing duration as well as a decrease in stride frequency for both models as compared to controls. These abnormalities persisted into adulthood in the *Nf1*^+/R681X^ model of NF1 but largely resolved in the CD model of WS, which displayed only one significant adult phenotype, a decreased percentage of the forelimb stride cycle spent in the stance phase compared to controls. The differences observed in spatial, temporal, and postural subcomponents of gait in both developing and adult *Nf1*^+/R681X^ animals suggest their gait abnormalities persist, while CD deficits largely disappeared by very early adulthood, with only select stride cycle features perturbed at this age. The CD model also displayed reduced intraindividual variability for several gait metrics during development, a feature that did not appear in adulthood or in the *Nf1*^+/R681X^ mice, suggesting this variability phenotype may be specific to the developing WS model and not necessarily to NDDs in general. Overall, the considerable overlap in abnormalities between these WS and NF1 models suggests that NDDs have multiple common gait phenotypes but may differ in their time course, with some abnormalities persisting into adulthood, while others reflect a developmental delay that eventually resolves.

Although the percentage of the stride cycle represented by swing or stance phases was the only metric significantly different from controls in the adult CD mice, several of the gait metrics perturbed during development have previously been reported to be altered in adult WS patient populations. While we observed significantly reduced intraindividual variability for several gait features in developing CD mice, increased gait variability has previously been documented in adults with WS, specifically for stride length and stance time as a percentage of the total gait cycle (Hocking et al., 2009). Stride length and hindlimb stance width were both altered in CD development, but while the mice showed increased stride length and reduced stance width, Hocking et al. (2008) reported that adults with WS exhibited reduced stride length and an increased base of support, equivalent to stance width (Krebs et al., 2002). This discrepancy in direction of effect may be attributable to the body length adjustment we included in our statistical model, which was not accounted for in the human study, or the fact that their assay recorded gait at a self-determined speed instead of a forced speed like DigiGait. Age differences likely also play a role, given the significant evolution of gait across time. Adult CD mice no longer showed the stride length and stance width abnormalities present in development, so it is possible that characterization of gait in older CD adults might reveal a continued trend of decreasing stride length and increasing stance width. Despite this apparent difference in direction of effect, the fact that similar gait metrics showed alterations in both human and mouse studies is interesting and should be explored in future studies, particularly with older mouse models and child and adolescent WS populations.

These data provide a foundation for investigating which genes in the WS critical region, in isolation or in combination, may be driving the gait abnormalities observed here. The transcription factors *Gtf2i* and *Gtf2ird1*, for instance, have been ruled out for most other WS neurobehavioral phenotypes (Kopp et al., 2019; Kopp et al., 2020), but may still be involved in the motor dysfunction. Future studies involving intercrossing of mutants harboring loss of function mutations in single WS genes will help to clarify the role of each of the WS locus genes in gait function.

The developmental gait differences between *Nf1*^+/R681X^ mice and controls also relate to findings previously reported in the human NF1 literature. The significantly increased stride length that we observed in developing *Nf1*^+/R681X^ mice, although not in the same direction of effect, relates to a previous study of children with NF1 which showed significantly decreased stride length (Champion et al., 2014). The increase in hindlimb stance width compared to controls that we observed across development in *Nf1*^+/R681X^ mice also differs from the decreased base of support reported in NF1 children compared to reference values. However, as noted above, these human studies did not employ a treadmill-based assay nor account for body size, both of which are known to affect numerous gait metrics (Akula et al., 2020; Herbin et al., 2007). Additional data collection in NF1 patient populations that standardizes walking speed by utilizing a treadmill or similar device will allow for more meaningful comparison to our Digigait forced-gait findings. Because few human NF1 gait studies have been performed, collection of more data across the lifespan should elucidate whether the spatial, temporal, and postural gait phenotypes we observed in *Nf1*^+/R681X^ mice reflect a similarly persistent phenotype in the patient population.

Crucially, the use of a patient-derived mutation in our *Nf1* model represents an increasingly feasible precision medicine approach to disease modeling, setting an example for how mouse models can be used in future studies of NDD mechanisms and interventions. There are multiple other genetically-engineered mouse models of *Nf1* mutations, both artificial (Brannan et al., 1994; Costa et al., 2002; Jacks et al., 1994; Maloney et al., 2018) and patient-derived mutations (Guo et al., 2019; Li et al., 2016; Toonen et al., 2016), which can be similarly assessed for gait function and compared to the present findings. Identification of the similarities and differences in gait phenotype across different *Nf1* models will allow for better identification of and interventions for the various motor deficits seen in NF1 clinical populations. In addition, we can use these patient-derived models to parse genotype-phenotype relationships in gait function as has been identified for other NF1 phenotypes such as optic glioma presentation and ASD features (Anastasaki et al., 2017; Morris & Gutmann, 2018).

The presence of gait abnormalities and the persistence of some into adulthood are also consistent with studies of other NDD mouse models. The *Mecp2* mouse model of Rett Syndrome displayed a reduced stride length in adulthood (Vogel Ciernia et al., 2017), one of the gait metrics we observed to be perturbed in both CD and *Nf1*^+/R681X^ models. Similarly, the valproic acid mouse model of ASD has previously been reported to exhibit a decreased stride length in the juvenile period compared to controls (Al Sagheer et al., 2018). Although this direction of effect is opposite the increased stride length we observed in both CD and *Nf1*^+/R681X^ developing mice, a correction for body length, such as we performed, might reveal greater similarity across models. Alternatively, the developmental trajectory of gait differences that we observed in *Nf1*^+/R681X^ mice may vary in timing between models, with some showing more prolonged developmental delays than others. While some metrics such as stride length are consistently characterized across NDD gait studies, many of the other variables examined here vary greatly between studies. A standard and comprehensive list of gait metrics to include in future studies would also benefit the field greatly by allowing more between-model comparisons.

Several limitations of rodent model gait analysis should be noted and provoke further studies of NDD-related gait phenotypes. Mice are quadrupeds instead of bipeds, and therefore extrapolation of mouse model gait results to human populations must be done cautiously. However, in our previous study of wild type mice, we documented gait differences across time that mirrored those previously observed in human gait development (Akula et al., 2020), and therefore we feel that the present study also provides meaningful information about the influence of WS locus hemizygosity and *Nf1* mutation on gait neurocircuit function. A substantial advantage of the DigiGait system is that it allows standardization of gait speed across mice, which is important because speed significantly contributes to gait variability (Jordan et al., 2007; Möckel et al., 2003); however, assays of spontaneous gait such as the CatWalk (Noldus) may allow for parsing of additional gait differences that reflect motivation-related behavior.

The gait phenotypes we have described here provide the basis for numerous future studies of NDD behavioral trajectories and suggest a potential outcome variable in the testing of NDD treatments. The concordance between gait phenotypes in our two NDD models is striking and unusual, as the establishment of phenotypes common to multiple NDDs has traditionally been a major challenge in the field when assessing more cognitive behaviors in rodent models. We have reported here a common signature between two NDDs in gait, a behavioral measure that is accessible and linked to a relatively well-understood circuit, and this methodology can be easily applied to other NDD mouse models to assess whether the observed gait abnormalities apply to a larger number of NDDs. If the gait phenotype observed here is common to other NDDs, that would suggest that this phenotype, even though not as central to the human definition of NDDs, might be a very sensitive and consistent gauge to assess what an NDD mutation does to a functioning CNS circuit. Investigating the related cellular and molecular consequences is also facilitated by the deep understanding of spinal motor circuitry relative to the more abstract circuitry characteristic of other NDD cognitive phenotypes. Finally, regardless of the presence or absence of commonalities, within a given NDD, gait could be widely used to test treatments across development, measure normalization of circuit function, and assess the effect of long-term therapies.

## Acknowledgments & Conflict of Interest Statement

The authors thank the Co-Directors of the Animal Behavior Core at Washington University School of Medicine, David Wozniak, PhD, and Carla Yuede, PhD, for access to the DigiGait equipment. Support for this study was provided by the NIMH (R01MH107515, JDD), NINDS (F31NS110222, RMR; R35NS097211, DHG), and NICHD (U54HD087011, Intellectual and Developmental Disabilities Research Center at Washington University). The authors declare that the research was conducted in the absence of any commercial or financial relationships that could be construed as a potential conflict of interest.

## Notes

### Competing Interest Statement

The authors have declared no competing interest.

